# Enhanced Immunoprecipitation Techniques for the Identification of RNA Binding Protein Partners: CRD-BP interactions in mammary epithelial cells

**DOI:** 10.1101/2021.06.10.447893

**Authors:** Saja A. Fakhraldeen, Scott M. Berry, David J. Beebe, Avtar Roopra, Vladimir S. Spiegelman, Caroline M. Alexander

**Affiliations:** McArdle Laboratory for Cancer Research, Pennsylvania State University College of Medicine; Department of Biomedical Engineering, Pennsylvania State University College of Medicine; Department of Neuroscience, University of Wisconsin-Madison and Pennsylvania State University College of Medicine; Department of Pediatrics, Division of Pediatric Hematology/Oncology, Pennsylvania State University College of Medicine

**Keywords:** Ribonucleoprotein immunoprecipitation (RIP), CRD-BP/IMP1/VICKZ1, exclusion-based sample preparation (ESP), PAR-CLIP

## Abstract

RNA binding proteins (RBPs) regulate expression of large cohorts of RNA species to affect programmatic changes in cellular phenotypes. In order to describe the function of RBPs within a cell, it is key to identify their mRNA binding partners. This is often done by cross-linking nucleic acids to RBPs, followed by chemical release of the nucleic acid fragments for analysis. However, this methodology is lengthy, involves complex processing leading to extraordinary losses, requires large amounts of starting materials, and is prone to artifacts due to the labile nature of mRNA. To evaluate potential alternative technologies, we tested “exclusion-based” purification of immunoprecipitates (oil-based IFAST™ or air-based SLIDE™), and report here that these methods can efficiently, rapidly and specifically isolate RBP-RNA complexes with minimal handling. The analysis starts with >100x less material than for techniques that include cross-linking. Depending on the specific antibody used, 50-100% of starting protein is retrieved, allowing the assay of endogenous levels of RBP instead of tagged and over-expressed ectopic proteins. Isolated protein and nucleic acid components are purified and analyzed using standard techniques to provide a comprehensive portrait of RBP complexes. Using exclusion-based techniques, we show that the mRNA binding partners for CRD-BP/IMP1/IGF2BP1/ZBP1 in cultured mammary epithelial cells are enriched in mRNAs important for de-toxifying superoxides (glutathione metabolic enzymes) and other mRNAs encoding mitochondrial proteins.

## 1. Introduction

RNA binding proteins (**RBP**s) are critical post-transcriptional regulators of gene expression in normal and pathological cellular contexts (1). At least 1542 RBPs govern RNA metabolism at myriad stages of splicing, export, storage, transport and translation (2). Often, RBPs bind select RNA species to modulate their expression, localization and/or stability, occasionally via highly specific and conserved sequence motifs. However, more typically, RBPs bind RNA species via short and degenerate sequences that are not easy to recognize prospectively (1).

Aberrant RBP activity is responsible for such important phenotypes as Fragile X syndrome (via the RBP FMRP) (3), and splicing reactions of cancer-associated tumor drivers, such as AR (via the RBP DDX3) (4). It is therefore important to define the cohort of mRNA binding partners that are bound by each specific RBP, since these are likely to be affected by altered RBP expression or activity. The cohort of mRNA species bound by a given RBP can be highly cell type-specific, for reasons that are not yet understood. RBPs sometimes stabilize mRNA species; this is deduced from the demonstration of a direct binding interaction, together with decreased abundance upon RBP knockdown/knockout (1, 5). However, other regulatory activities that do not result in altered RNA abundance are much more difficult to identify, for example regulation of RNA localization and delivery of target proteins to specific subcellular structures (6). It may be possible to evaluate these partnerships from a catalog of specific mRNA binding partners, for example the adhesion defects observed by Conway et al (7) for CRD-BP/IMP1 knockdown embryonic stem cells.

Most studies rely upon UV-induced cross-linking coupled with immunoprecipitation techniques to define mRNA binding partners for RBPs. By exploiting the unique chemical reactivity of RNA for protein, irreversible cross-links can be formed between RNA and protein moieties that lie in close proximity (1). This technique was widely adopted after concerns were raised about the potential for switching of RBP binding partners during incubations (8). However, it is not trivial to reverse these cross-links sufficiently to release and identify the bound mRNA species, and the yield of input RBP that emerges after the extensive processing reactions is substantially less than 1% of total (9). Various versions of photoactivatable-ribonucleoside-enhanced crosslinking and immunoprecipitation (**PAR-CLIP**) protocols have been described and applied to the analysis of RBPs; the strengths and weaknesses of each have been reviewed (5, 9–12). In general, the limitations of cross-linking protocols fall into various classes: the loss of unstable RNA species during long processing procedures, loss of mRNAs with indirect or low affinity interactions during washing of immunoprecipitates, artifacts created by cross-linking and extensive derivatization processes, and the requirement for an impractically high starting numbers of labeled cultured cells over-expressing the RBP of interest. Indeed, in many key cell types, RBP interactions cannot be studied due to the requirement for up to 1g of starting protein lysate. Here, we have applied two exclusion-based sample preparation (ESP™) technologies to identify mRNA binding partners for an exemplar RBP; the simple, expedited processing that is required for the techniques used in this paper versus those required for PAR-CLIP are summarized in **Fig. 1**.

**Figure 1.**
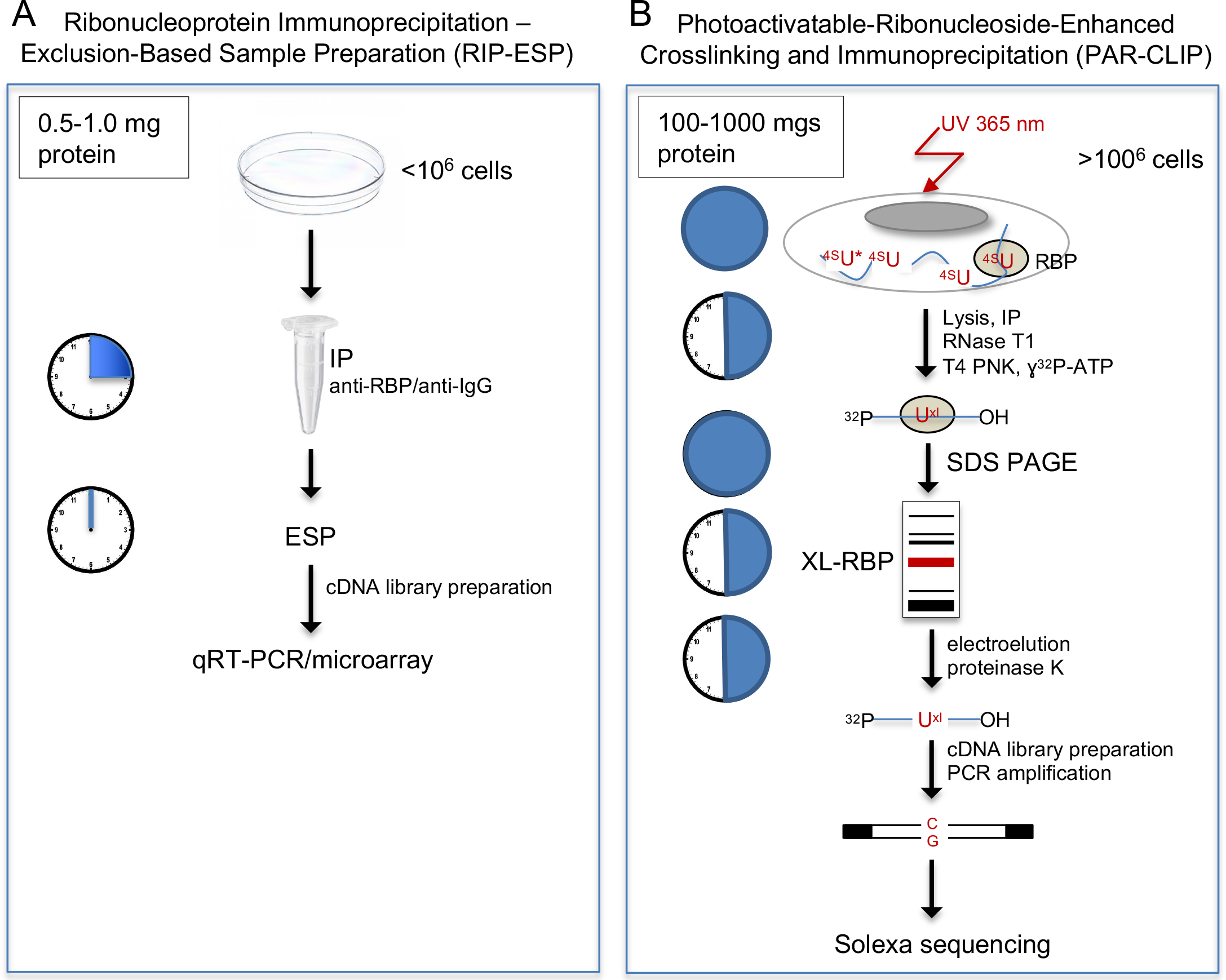
Comparison of ESP and (PAR-)CLIP technologies. Schematics of the workflow for ESP methodology (A) compared to PAR-CLIP (B), as performed by Hafner et al (37), to illustrate one of the incentives for performing this study. Specifically, the timeline and handling required for both procedures is shown on clock faces, together with the relative amounts of starting cell lysate required, and the complexity of the PAR-CLIP protocol. Typically, RBPs are over-expressed for CLIP protocols. * indicates the use of nucleoside substitution, which can induce a nucleolar stress response and result in cytotoxicity (71, 72).

The focus of this study is the RNA binding protein <u>C</u>oding-<u>R</u>egion <u>D</u>eterminant <u>B</u>inding <u>P</u>rotein (CRD-BP), originally defined as a regulator of stability of Myc mRNA. CRD-BP has been independently identified as to protein responsible for several important activities, as reflected by its diverse set of names (including IMP1, IGF2 mRNA binding protein, gene name is IGF2BP1; ZBP1, zipcode binding protein-1, VICKZ1) (13–16). This protein is often highly expressed in tumor and fetal cells and is required for several tumor-associated activities in a wide range of cell types (17–21). We showed previously that both normal breast epithelial cells and breast tumor cells expressed relatively low levels of two forms of the CRD-BP protein; despite low expression, this protein regulated clonogenic growth of breast cancer cells *in vitro* (22). Indeed, CRD-BP has been shown to be required for clonogenic activity in several tumor cell types, suggesting that it enables some fundamental property required for clonogenic growth (23).

Target mRNAs bind the KH repeat domains of CRD-BP via combinatorial interactions through a looped tertiary structure with short consensus sequences. This complex interaction makes them difficult to predict *a priori* (24, 25). To understand the molecular basis of important functional activities, we tested whether ESP technologies could isolate mRNA binding partners from the low levels of CRD-BP that are expressed by breast epithelial cells. This focused our attention on efficient retrieval of CRD-BP complexes. Exclusion based technologies have been deployed for many reasons, including their sensitivity, speed, parallel processing capacity and potential for multiple endpoint assays (26). The mRNA binding partners identified by this analysis included the mRNAs for glutathione metabolism, including the seleno-protein glutathione peroxidases Gpx1 and Gpx2 (important for the detoxification of superoxides (27)), and a group of mRNAs encoding proteins destined for mitochondria. We propose that this technology is a useful approach to dissecting RBP function, either alone, or as a complement to techniques requiring accurate binding site predictions that are typically derived from cross-linking studies.

## 2. Materials and Methods

### 2.1 Cell culture and transfections

Mouse embryonic fibroblasts (MEFs), the EP and EN sub-strains of the mouse mammary epithelial cell line HC11, 293T, MCF7 and MDA-MB-231 cells were cultured as previously described (22). A construct expressing Flag-tagged CRD-BP was described in a previous publication (28). Transfections were carried out using Lipofectamine LTX with plus reagent (Life Technologies) according to manufacturer’s instructions.

### 2.2 Ribonucleoprotein immunoprecipitation (*RIP*)

Cells were cultured in 10 cm dishes and lysed in polysome lysis buffer (10 mM HEPES (pH 7.0), 5 mM MgCl_2_, 100 mM KCl, 0.5% NP-40) with freshly added DTT (1 mM), RNase (100 U/ml), and protease/phosphatase inhibitors (Thermo Fisher Scientific). Approximately 10^6^ 293T cells (a single 10 cm dish at ~80-90% confluence) yields 500 μl of lysate, with approximate protein concentration 4 μg/μl; RIP reactions were processed in batches of 200 μl. For other cell types, such as the breast epithelial cells described in these studies, 2× 10cm dishes were processed into 500 μl of lysis buffer. Lysates were sonicated (10 pulses at 4-5W) and cleared by spinning 3× at 12,000 rpm for 30 minutes at 4°C. Protein concentrations for the whole cell lysates and the unbound fractions were determined using Bradford reagent (Sigma-Aldrich). For immunoprecipitation, approximately 1.2 μgs of specific primary antibody (or matched IgG control) were added, together with 5 μl of washed protein G-bound paramagnetic Dynabeads (Life Technologies cat#10003D). Antibody binding of RBP complexes was allowed to proceed at 4°C for the times indicated, before purification using IFAST, SLIDE-based or standard immunoprecipitation. The following antibodies were used: anti-CRD-BP antibody (Cell Signaling cat#8482; RRID:AB_11179079), anti-Flag antibody (Sigma-Aldrich cat#F3165; RRID:AB_259529), or non-immune rabbit IgG control (Jackson Immunoresearch cat# 011-000-003; RRID:AB_2337118). Following the RIP procedures, aliquots from the bound and unbound fractions were harvested and assayed for CRD-BP using Western blotting (to assess efficiency of extraction), and the remaining sample was used for RNA analysis.

### 2.3 Western blotting

Lysates were analyzed by SDS-PAGE followed by transfer to PVDF membranes, as described in (22). Primary antibodies: anti-CRD-BP (Cell Signaling cat#8482; RRID:AB_11179079), diluted 1:1000; anti-vinculin (Millipore cat#05-386; RRID:AB_309711) diluted 1:5000; anti-GAPDH (Cell Signaling cat#2118; RRID:AB_561053) diluted 1:3000-1:5000. Secondary antibodies: HRP anti-mouse (Jackson Immunoresearch cat# 715-035-151; RRID:AB_2340771); HRP anti-rabbit (Invitrogen cat# G-21234 RRID:AB_2536530) diluted 1:5000 or HRP mouse anti-rabbit IgG (conformation-specific) (Cell Signaling cat#L27A9; RRID:AB_10892860). All antibodies were diluted in 5% milk in TBS-Tween.

### 2.4 RNA isolation and analysis

Total RNA was isolated using the RNeasy Mini Kit (Qiagen). RNA concentrations and quality were determined using a Nanodrop instrument (Thermo Scientific; average 260/280 ratio ~2.1). Reverse transcription was performed as previously described, and specific mRNAs were quantified using q-RT PCR (29).

### 2.5 Microarray analysis

A Nimblegen 12-plex whole mouse genome microarray chip (Build 100718_MM9_EXP_HX12; comprising 44171 probes, equivalent to 24205 individual genes) was used to assay the relative abundance of each mRNA in cDNA libraries made from each sample. RNAs were processed for this analysis according to manufacturer’s instructions; briefly, first strand followed by second strand cDNA synthesis was performed, followed by RNase cleanup, and cDNA precipitation. Double stranded cDNA (4 μg) was Cy3-labeled and hybridized to microarray chips, which were then washed and scanned. For any given experimental condition, duplicate sets of four samples of cDNA were prepared for analysis by microarray: unbound and bound RNA from anti-CRD-BP immunoprecipitates, and from anti-IgG immunoprecipitates (specificity control) for analysis. The data was analyzed using Multi-Experiment Viewer (MeV) software.

### 2.6 Bioinformatic analysis

Raw intensity readings from the microarray analysis were log2-transformed and median-centered. These lists were rank-ordered, and the rank of genes determined for the mRNAs co-purifying with the paramagnetic particles (**PMP**s), for comparison with the ranking of genes in the unbound fractions. Statistical analysis of independent replicates was used to reflect the significance of fold changes, p <0.01. Enriched gene sets for RNAs pulled through by the specific antibody, anti-CRD-BP, were compared with the non-specific control, IgG fraction. The relative enrichment of RNA species in bound fractions was confirmed independently by qRT-PCR. These confirmed mRNA species, accumulating in anti-CRD-BP immunoprecipitation reactions, were imported into pattern prediction algorithms, such as STRING (string-db.org).

## 3. Results

### 3.1 Use of exclusion-based sample preparation (*ESP*) devices

**IFAST** (<u>I</u>mmiscible <u>F</u>iltration <u>A</u>ssisted by <u>S</u>urface <u>T</u>ension): IFAST devices are fabricated from polypropylene via injection compression molding (DTE Research and Design, LLC), and consist of linearly aligned wells (5-15 μl volume) connected by trapezoidal microfluidic conduits (**Fig. 2**). These wells are flanked by a larger input well (up to 200 μl volume) on one end, and an output well of designer-specified volume (5–10 μl) on the other. The pre-incubated cell lysate/antibody/paramagnetic particles (**PMP**s; prepared as described in Materials and Methods) mixture is transferred to the input well, and PMP-bound biomolecular complexes are purified by a magnet-based pull through the intermediate wells, which consist of alternating solutions of oil (Fluoinert FC-40 oil, Sigma-Aldrich), and aqueous wash phases, to the output well. Note that this requires no pipetting or additional handling beyond the initial loading of the device and takes an average of 20-30 seconds. The utility of this device for identifying valid biological interactions (including weak interactions), for streamlining multiplexed assays of analytes, and for the detection of viral RNAs for clinical diagnostics has been previously demonstrated (30–34).

**Figure 2.**
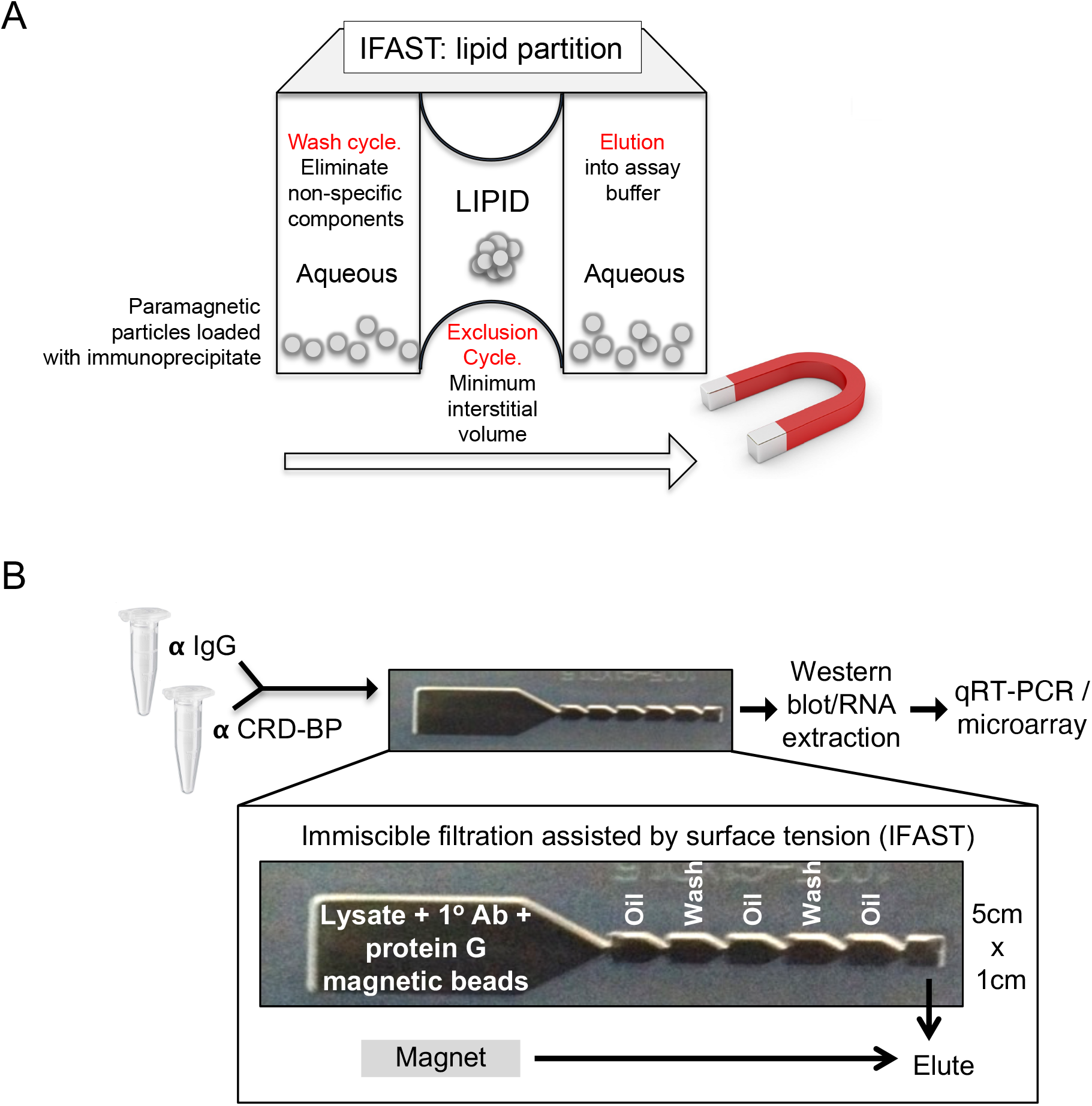
Overview of ESP-based Immiscible Filtration Assisted by Surface Tension (IFAST). **A.** Schematic of RIP-IFAST technique; the purification (or “exclusion”) phase of the ribonucleoprotein immunoprecipitation process is provided by pulling magnetic beads loaded with immunoprecipitate through a lipid barrier located between aqueous wells. **B.** An overview of the IFAST protocol, showing the configuration and dimensions of the device.

**SLIDE** (<u>S</u>liding <u>L</u>id for <u>I</u>mmobilized <u>D</u>roplet <u>E</u>xtractions): In order to avoid the use of oil-based exclusion, we employed a SLIDE device, which depends instead on air-based exclusion (35). This has the advantage of eliminating oil from the purification process and the pull-through lysate. The SLIDE device consists of a handle and a base, each with movable magnets within them (35) (commercial name is EXTRACTMAN™ from Gilson). A polypropylene well plate (provided by Gilson, Inc.) is loaded with samples containing PMPs, wash buffer, and elution buffer. By sliding the SLIDE handle across the base, PMPs are rapidly and efficiently transferred between reagents in series. Importantly, PMPs are collected on a disposable PMP collection strip, which is comprised of highly polished uncharged polypropylene. Thus, this hydrophobic PMPs collection minimizes carryover of aqueous material as the SLIDE handle moves between reagents. In RIP experiments, the input wells of this device are loaded with cell lysate and the RBP-complex bound-PMP beads are moved through adjacent wells containing wash buffer as described above (**Fig. 3**); total time for exclusion purification is approximately 20 seconds.

**Figure 3.**
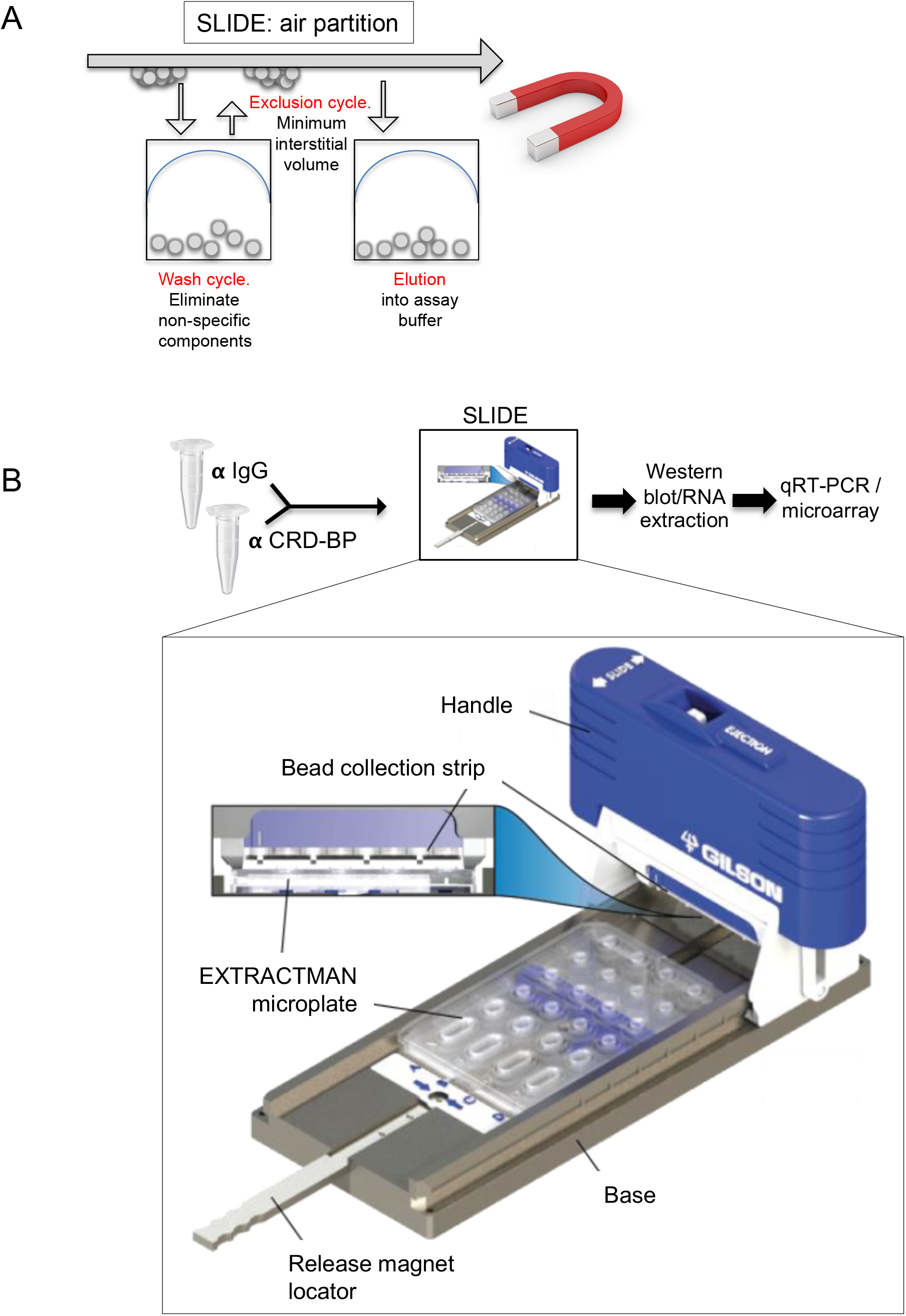
Overview of ESP-based Sliding Lid for Immobilized Droplet Extraction (SLIDE). **A**. Schematic of SLIDE™ technique using EXTRACTMAN™ device (Gilson, Inc.). The exclusion phase operates by repeated cycles of lifting of magnetic beads loaded with immunoprecipitate out of the wash solution. The extracted drops are held against a hydrophobic surface with minimum aqueous volume and surface tension prior to re-elution in a fresh aqueous solution. **B**. An overview of the SLIDE protocol, showing the configuration and dimensions of this specific device.

By passing the PMPs carrying immunoprecipitation complexes through oil (IFAST) or air (SLIDE) by attraction to a magnet, the aqueous dead volumes are minimized, reducing the time and handling required to dilute out associated fluids (i.e. to wash immunoprecipitates). The internal aqueous volume of paramagnetic particles (PMPs) is approximately 115 nls for each 5 μl volume of beads. We optimized the protocol for this specific buffer composition, given that the residual surface volume determines surface tension (increased by higher salt and decreased by detergent). Samples were processed simultaneously for up to four immuno-PMP lysates (directly in parallel), whereas samples were processed individually in the IFAST devices.

### 3.2 Efficiency of purification of the RNA binding protein, CRD-BP

We tested the efficiency of the recovery of endogenous CRD-BP protein by immunoprecipitation using IFAST, first for two cell types, 293T human embryonic kidney cells and mouse embryonic fibroblasts (MEFs), and then for a tagged CRD-BP protein (applying a different antibody, anti-FLAG) expressed in cultured mouse mammary epithelial cells (**Fig. 4A, B**) (36, 37). Using the anti-CRD-BP antibody to purify endogenous CRD-BP from 293T cells and MEFs, the yield of purified protein was 50-60% of total input; for the high affinity anti-FLAG antibody, losses were insignificant. Vinculin was used as an indicator of non-specific protein adsorption, and was not detectable. We also tested the efficiency of the affinity purification by assaying residual antibody in the unbound fraction and found almost no losses for the immunocomplexes during extraction from the cell lysates (**Fig. 4C**).

**Figure 4.**
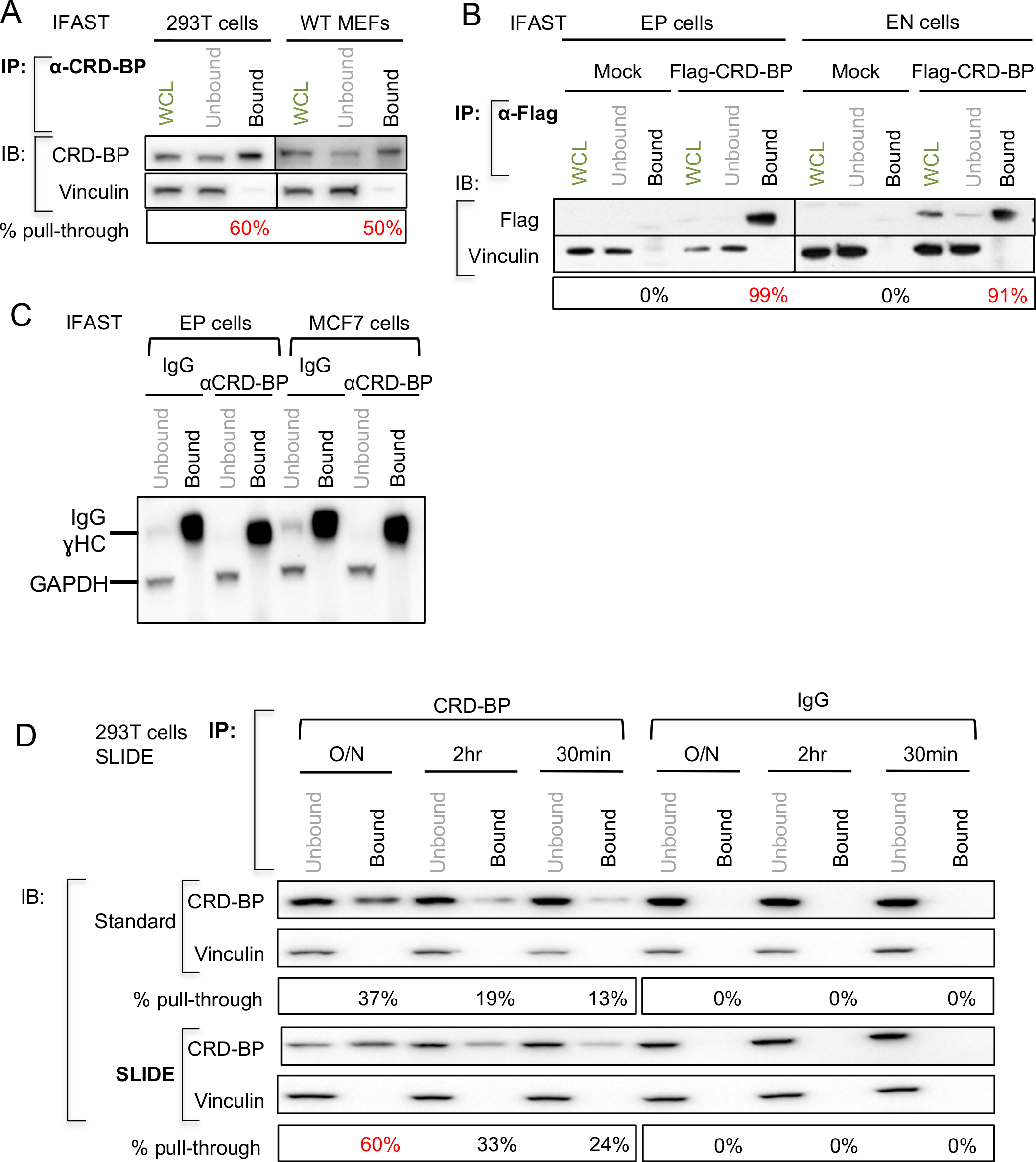
Efficiency of pull-through of CRD-BP protein using IFAST. **A**. *Demonstration of efficiency of pull-through of endogenous CRD-BP.* Two cell types with high levels of endogenous CRD-BP (293T and MEF) were lysed, incubated with anti-CRD-BP primary antibody and protein G paramagnetic particles (PMPs), and RIP fractions purified using IFAST. A known amount (typically 20%) of total immunoprecipitate pulled through was analyzed by Western blotting (bound) and compared to the input remaining (unbound), with <u>w</u>hole <u>c</u>ell lysate (before pull-through; **WCL**) shown for comparison. Vinculin is used to evaluate specificity of the immunoprecipitation. The fraction of CRD-BP pulled through is shown below (%). **B.** *Comparison of efficiency of pull-through of tagged CRD-BP.* Mouse mammary cell lines, EP and EN cells, were transfected with Flag-tagged CRD-BP (or empty vector, **mock**). 48 hours later, lysates were purified using IFAST. **C**. *Evaluation of efficiency of pull-through of antibody-PMP particles using IFAST*. For the cell lysates indicated, the amount of IgG remaining in the unbound fraction was assessed by Western blotting with conformation-sensitive anti-IgG antibody. **D.** *Evaluation of efficiency of immunoprecipitation with time*. 293T cell lysates were incubated with anti-CRD-BP antibody or an IgG control and paramagnetic particles for varying lengths of time (overnight (O/N), 2 hours, or 30 minutes) prior to purification either by standard or SLIDE-based RIP. Pull-through efficiency was evaluated by Western blotting.

We next evaluated the efficiency of recovery when shorter times were allowed for immune-complexation. Maximal recovery was found for overnight incubation, but significant recovery was obtained using only 30 minutes of binding (24% for 30 minutes compared to 60% recovery for overnight complexation) (**Fig. 4D**). For unstable RNAs, these short preincubation times could be particularly important.

### 3.3 Efficiency of immunoprecipitation of RNA with endogenous CRD-BP protein

To evaluate the efficiency of recovery for cells with low endogenous levels of CRD-BP, we tested mouse mammary epithelial cells (EP cells). Although CRD-BP is typically 100x less expressed in cells derived from adults compared to fetal cells, CRD-BP is still functionally important, at least for the expression of clonogenicity *in vitro* (22). We showed that the efficiency of pull-through of CRD-BP by IFAST from mouse mammary epithelial cells (EP cells) was approximately the same as for the cell lines with high endogenous levels of CRD-BP (shown in Fig. 4), measured at 62% by Western blotting (**Fig. 5A**).

**Figure 5.**
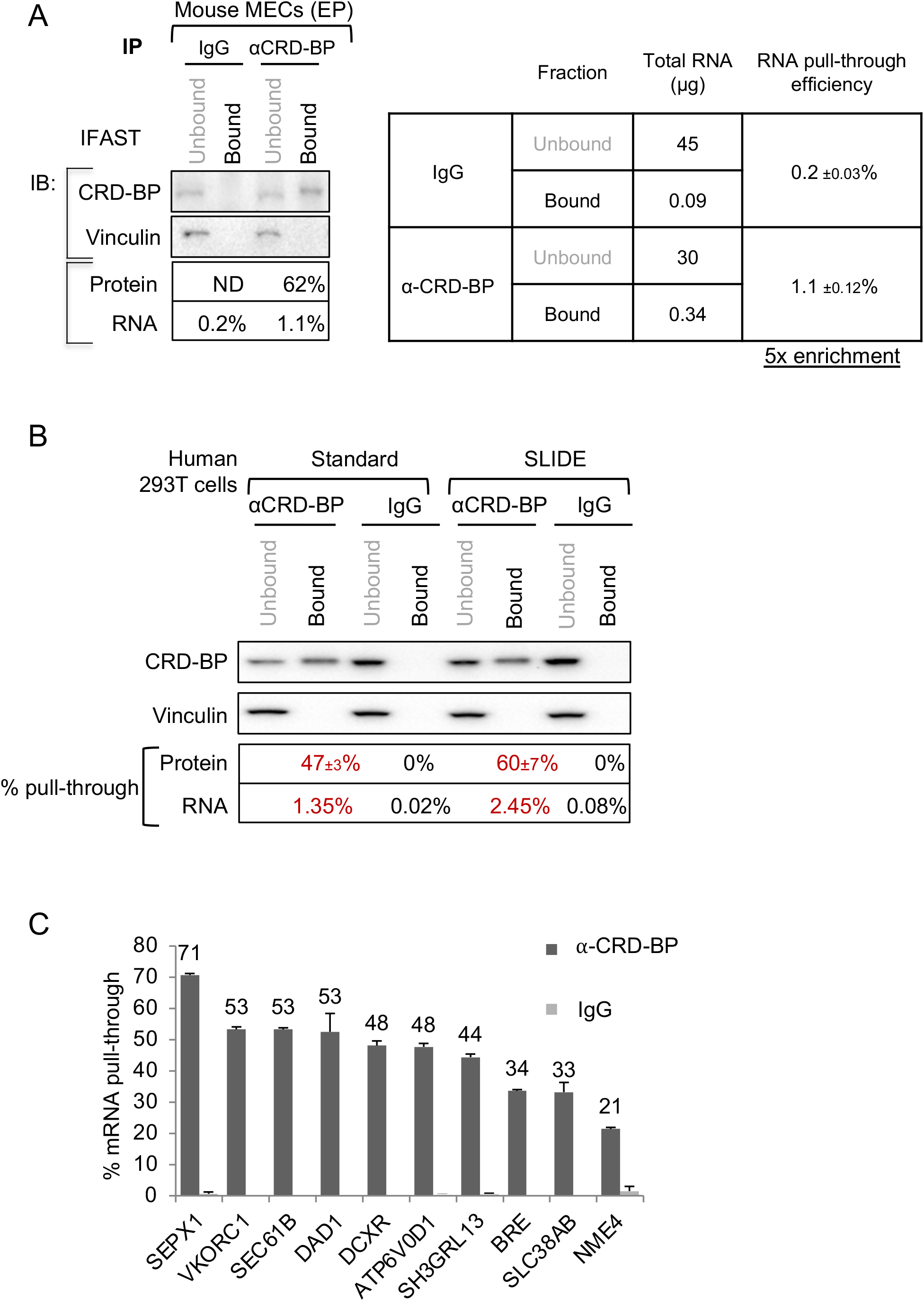
Immunoprecipitation of RNA in RIP CRD-BP complexes isolated by ESP methods. **A.** *Determination of amount of RNA in IFAST RIP.* Mouse mammary epithelial (EP) cell lysates were incubated with anti-CRD-BP antibody or an IgG control, and immunoprecipitates purified using RIP-IFAST. The protein component of the immunoprecipitate was analyzed by Western blotting as for Figure 4. RNA was purified and the amount of RNA in each fraction was determined. **B.** *Side-by-side comparison of RNA purification by standard and SLIDE RIP*. 293T cell lysates were incubated with anti CRD-BP 1° Ab or an IgG control, and lysate + antibody mixtures were purified using either standard or SLIDE-based RIP. The efficiency of pull-through of CRD-BP (n=3) and associated RNAs (n=2) is shown. **C.** *Validation of SLIDE-enriched mRNA partners.* Selected RNA species identified in Flag-tagged CRD-BP-associated RNP particles by Jonson et al (36) were evaluated by qPCR of SLIDE-enriched RIP fractions of endogenous CRD-BP from 293T cells.

Using the IFAST protocol, 5-fold more RNA was pulled through with the CRD-BP immunocomplexes than with the control (IgG) bound PMPs (1.1% compared to 0.2% for anti-IgG) (**Fig. 5A**). A “standard” immunoprecipitation protocol without cross-linking was compared with IFAST-purified RIP complexes; in other words, we used typical serial pipetting operations to conduct sequential, manual washes of each immunoprecipitate-bound PMP sample in individual Eppendorf tubes. We found broadly similar efficiency for recovery of both RBP protein and the total associated RNA (**Fig. 5B**).

We also tested whether the RNAs pulled through by this enhanced immunoprecipitation protocol included mRNA binding partners previously characterized as CRD-BP/IMP1 binding partners in 293T cells (36). All ten mRNA species surveyed were significantly pulled through by SLIDE-RIP (**Fig. 5C**).

### 3.4 Analysis of the CRD-BP mRNA complexes from mouse mammary epithelial (EP) cells

Ranked gene lists of bound and unbound mRNA fractions were compared for CRD-BP and IgG control IFAST-purified RIP fractions, to identify species that showed a significant change in rank listing (p<0.01). The CRD-BP gene list included 1,343 genes, of which 443 (approximately 35%) overlapped with the gene list from IgG control fractions (1,170 genes). These “sticky mRNAs” were subtracted from the total to generate a list of 900 potentially specific mRNAs in CRD-BP-associated complexes.

The fold enrichment of these 900 mRNA species (all >2-fold) is illustrated in **Fig. 6A**, and the mRNAs most highly enriched are shown as **Fig. 6B** (>4-fold). To verify the array analysis of RIP fractions, we selected >30 mRNA species for evaluation by qRT-PCR, including enriched and excluded mRNAs (**Fig. 6C**). For the purpose of illustration, we set a threshold on this confirmation assay; this threshold excludes 93% of mRNAs not enriched by array analysis, and also increases the stringency of inclusion in the specifically enriched fraction.

**Figure 6.**
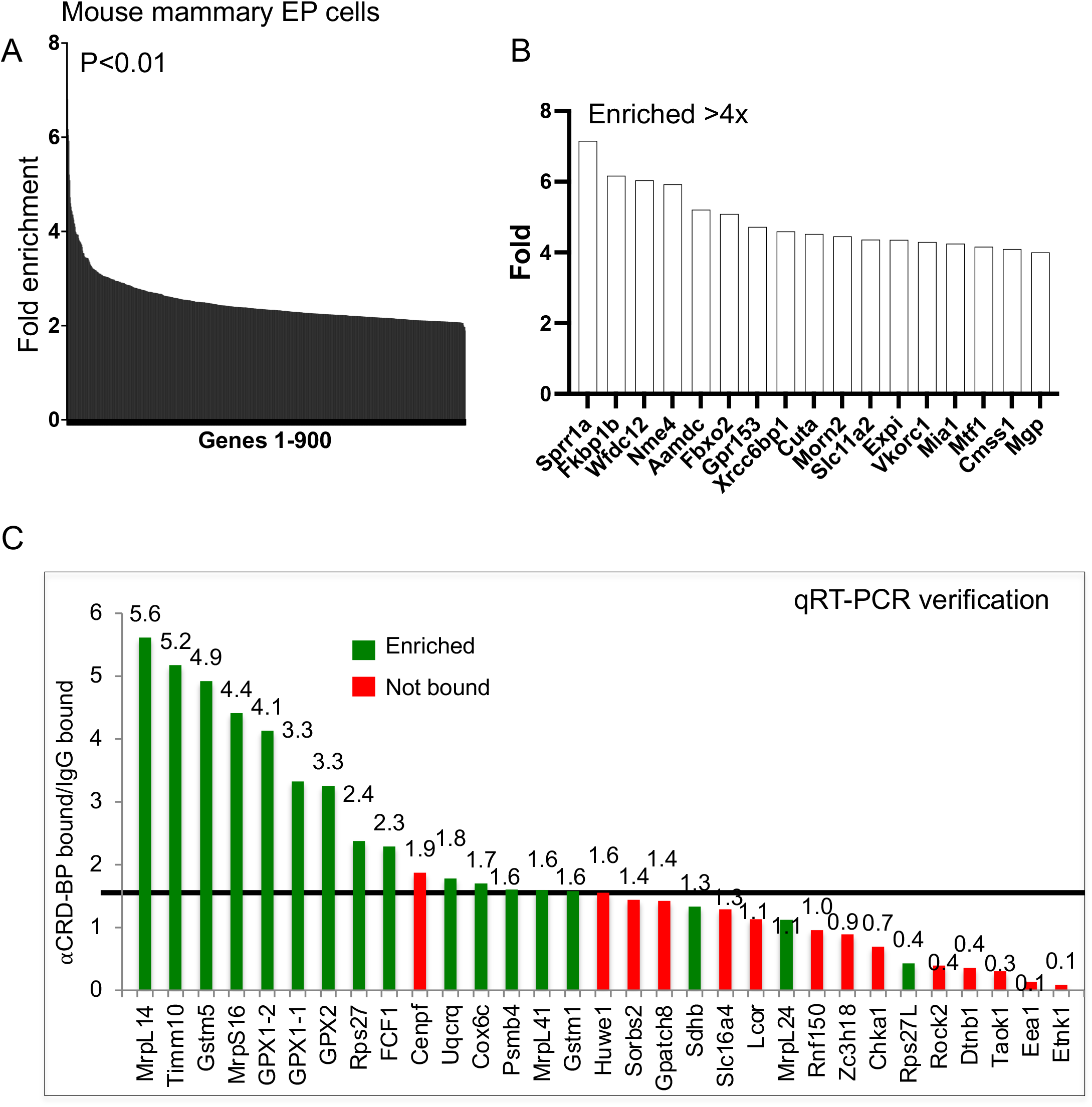
Analysis of RNA binding partners for CRD-BP in mouse mammary epithelial cells (EP cells). **A.** *General enrichment of bound mRNAs*. The relative fold-enrichment of 900 genes judged as specifically enriched in the anti-CRD-BP complexes (p<0.01) purified by IFAST-RIP from EP cells. **B.** *Most enriched mRNA species.* The mRNA species >4x enriched are shown as a detail of panel A. **C.** *Confirmation of mRNAs pulled through in functional groupings.* Functional groupings are shown diagrammatically in Fig. 7. To confirm enrichment scored from array analysis, a subset of associated mRNAs were tested by qRT-PCR analysis. For this study, a set of 31 mRNAs comprising 17 specifically and significantly enriched in the CRD-BP-bound fraction (green), were compared to 14 mRNAs specifically excluded from the CRD-BP-bound fraction (red). A potential thresholding line is drawn, that excludes 93% of mRNAs not enriched by array analysis and includes 64% of mRNAs designated as enriched.

To test whether this group of genes includes mRNAs associated with related cellular processes, we analyzed the group of 900 genes by STRING analysis (**Fig. 7**). We found significant enrichment of genes involved in glutathione metabolism, including glutathione peroxidase-2 (Gpx2), which catalyzes the reduction of organic hydroperoxides and H_2_O_2_ by glutathione, protecting cells against oxidative damage. Also enriched were glutathione S-transferases of the mu, theta and omega classes (Gstm1, −5, Gstt1 and Gsto2), involved in detoxification of electrophilic compounds (including carcinogens, therapeutic drugs, environmental toxins and products of oxidative stress by conjugation with glutathione microsomal glutathione-S-transferase), and microsomal glutathione S-transferase (Mgst3) involved in the production of leukotrienes and prostaglandin E. Another group includes core components of DNA synthesis. The other two groups enriched in this complex reflect mRNAs that are made in the nucleus and are translated proximal to the mitochondrion (38), including the mitochondrial ribosomal components (labeled mitochondrial translation) and complex 1-associated mRNAs of the electron transport chain that are not made by the mitochondrion (labeled Complex 1). The gene lists identified by these analyses are provided as supplemental data (Table S1).

**Figure 7.**
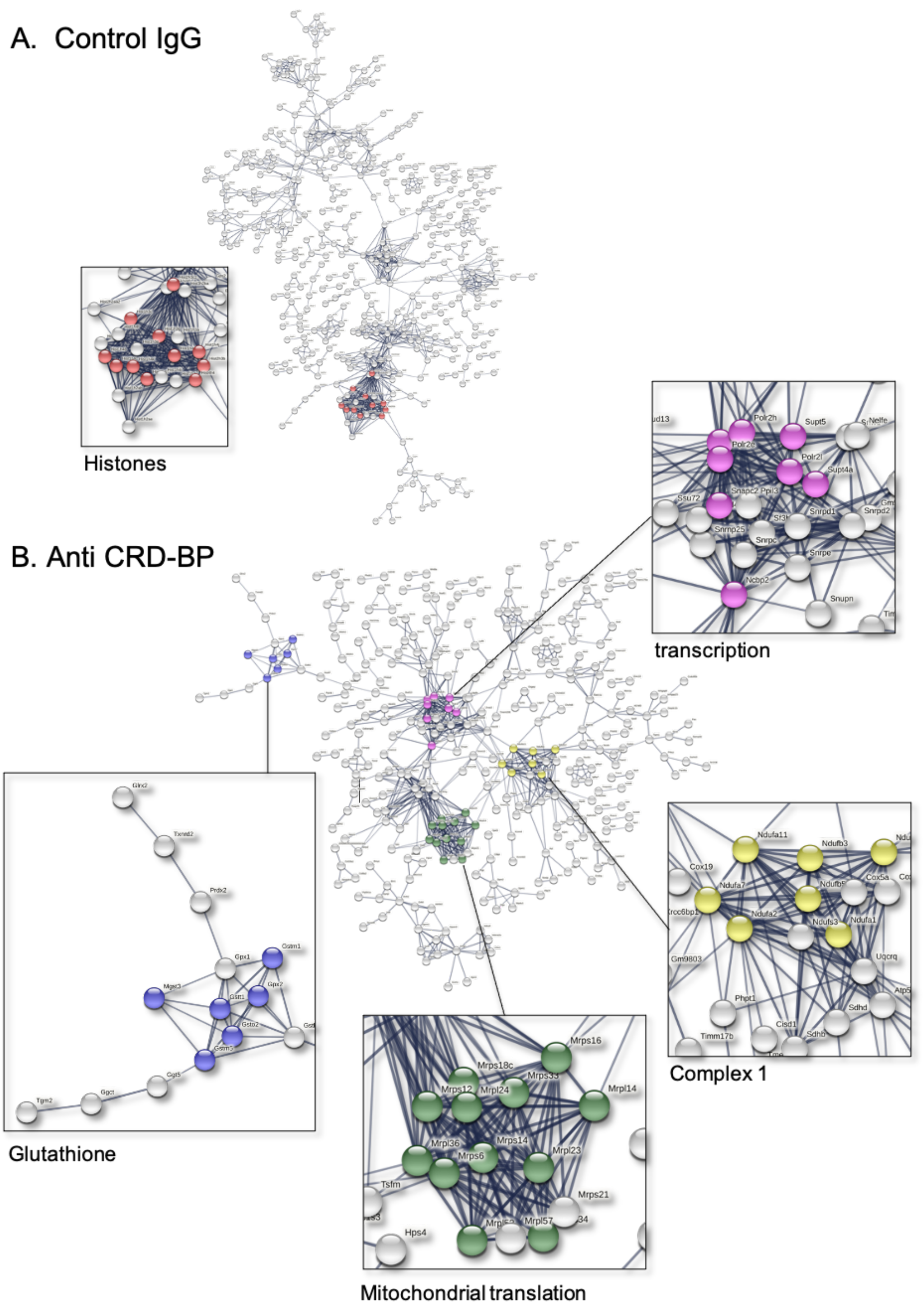
Functional grouping of mRNAs in CRD-BP complexes from mouse mammary epithelial (EP) cells. **A.** The mRNAs in control IgG complexes were subjected to STRING analysis, which found no significant enrichment groups except for a (relatively unique) group of histone mRNAs, and intriguingly, the mRNA for CRD-BP itself. **B.** The mRNAs in CRD-BP-associated complexes showed functional enrichment for 4 groups of genes, glutathione metabolism genes (including Gpx2), mRNAs related to transcription, and mRNAs for proteins targeted to mitochondria, important for mitochondrial ribosome function and the assembly of complex 1 in the electron transport chain.

### 3.5 Corresponding mRNAs were pulled through in CRD-BP-associated complexes from breast tumor cells

By way of preliminary validation of these results in human breast cancer cells, lysates of MCF7 and MDA-MB-231 cells were purified by anti-CRDBP RIP-SLIDE, showing a protein purification efficiency of approximately 47% for MCF7 cells, together with a yield of 2.6% mRNA. This included over 7-fold enrichment (CRD-BP/IgG) and >25% total yield of GPX1 (glutathione metabolism), MRPS16, MRPL14 and TIMM10 (mitochondrial targeted mRNAs), compared with <7% yield of excluded mRNA species (TAOK1 and ETNK1) (**Fig. 8**).

**Figure 8.**
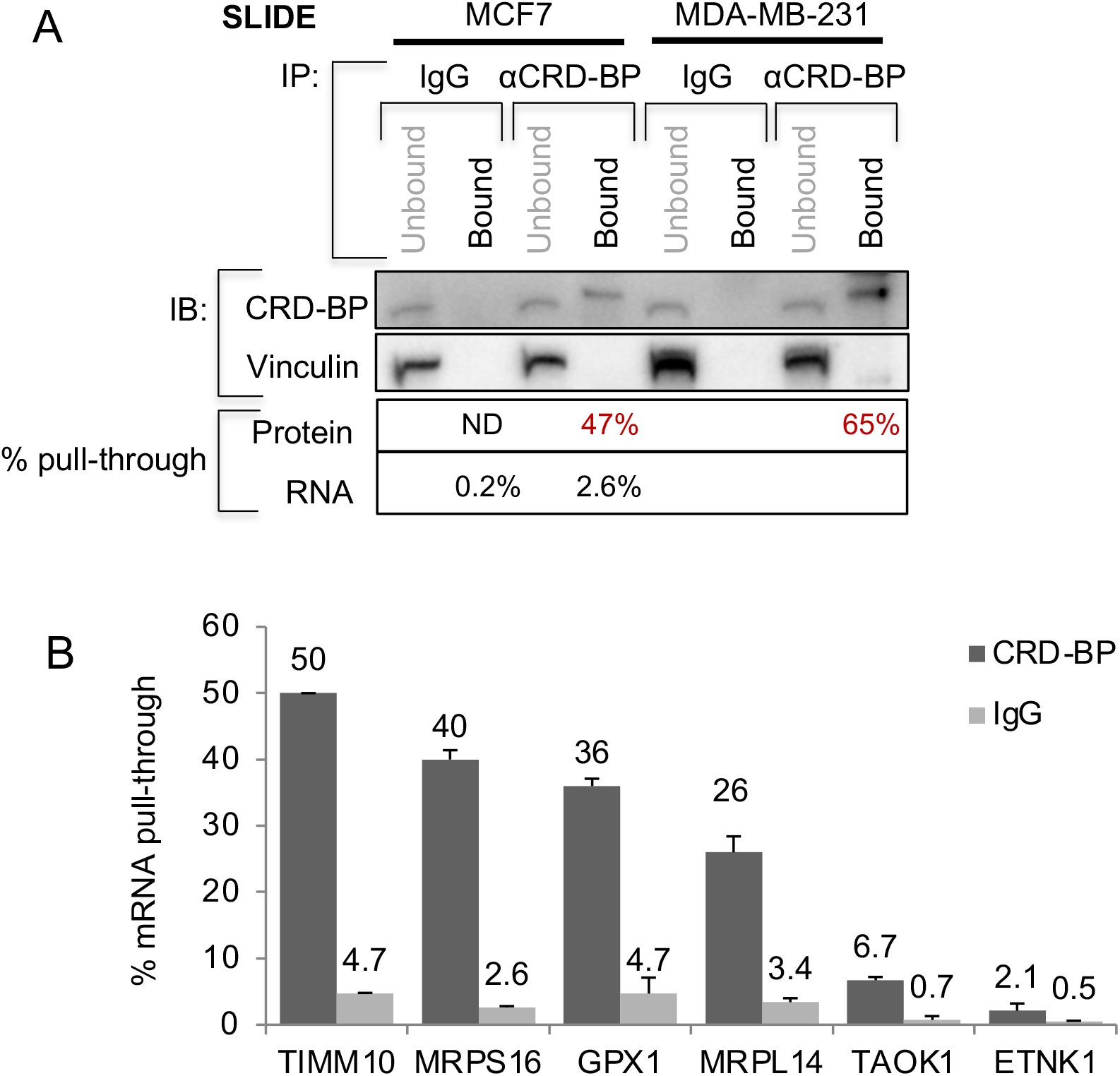
Confirmation of ROS-detoxifiers and mitochondrial mRNAs amongst CRD-BP binding targets in breast cancer cell lines. **A.** *Comparison of RNA targets enriched by SLIDE processing of CRD-BP RIP from breast cancer cell lines.* MCF7 and MDA-MB-231 human breast cancer cell lysates were incubated with anti-CRD-BP antibody or an IgG control, and the lysate + antibody mixtures were subjected to purification by SLIDE (as per Figure 4) and the efficiency of pull-through of CRD-BP protein and RNA binding partners was calculated. Six mRNAs were assayed by qRT-PCR, including four specifically associated in RIP fractions of mouse mammary epithelial cells, and two excluded from these fractions (TAOK1 and ETNK1; Fig. 6).

## 4. Discussion

### 4.1 The application of ESP to the discovery of RNA binding partners for proteins

Exclusion-based sample preparation (ESP) has been shown to be useful for cell, DNA and protein isolation from complex biological samples (35, 39, 40). The main time and labor-saving aspects of ESP methods are the substitution of the standard, manual pipetting operations with a swift, coordinated wash via magnetized bead adsorption. There can be profound impact of this technology, since increasing the speed of purification promotes the identification of more weakly bound or labile complex components (30). This may particularly apply to RNA-protein interactions, since the potential for exchange during the rapid wash procedure applied with ESP technologies is low. The potential for exchange of interactors was one of the drivers for development of cross-linking technologies to the study of RBP complexes. These concerns arose from a letter to the editor of *RNA* in 2004 which showed that 90 mins of co-incubation of a lysate containing an RNA binding protein (HuR) together with a cell lysate containing a known specific binding partner, the *fos* mRNA species, was more than enough to promote their interaction (8). We showed in a previous publication that dissociation of an antibody/protein complex increased from zero to 80% dissociation during a timecourse of 100 minutes (30). Only 10 minutes in a moderate salt wash buffer promoted 50% complex dissociation, but one minute showed insignificant losses, and importantly, the specificity of the wash was maintained. Therefore, the timescale for the ESP purification (on the order of seconds) prevents re-association artifacts.

### 4.2 Summary of advantages of ESP over RIP-ChIP and CLIP technologies

Techniques for identifying the RNAs that associate with specific RBPs have become increasingly sophisticated (10, 41, 42). However, the downsides of crosslinking of RNA to RBP have been noted before in a publication that showed optimal isolation conditions for a classic “RIP-ChIP” (43). A summary of the processing aspects of the ESP-based technologies, IFAST and SLIDE, versus one of the examples of CLIP technologies, PAR-CLIP (44–47), is shown in Figure 1.

We summarize these potential advantages of ESP:

i. *Almost quantitative yields.* The amount of pull-through of any given RBP is related to the avidity of the antibody for the protein of interest (shown by the almost quantitative extraction of a Flag-tagged version of CRD-BP by the high-avidity anti-Flag antibody, Fig. 4B). For the anti-CRD-BP antibody, the total yield is approximately 60% of total. Unlike previous RIP studies, we can claim to examine the binding interactions of the majority of RBP protein, instead of <1% of total (48). This also offers the opportunity to study endogenous level proteins, where the exogenous expression of RBPs can be mis-localized (36).
ii. *Starting amounts of lysate*. The amount of starting material required is over 100× less than for a typical CLIP procedure, given there are virtually no losses. Indeed, the sensitivity could be enhanced still further, depending upon the output required. This is especially important for human samples and samples of purified or limiting cells of any source (such as subpopulations of tumor cells or stem cell populations).
iii. *No mechanistic bias is required.* The underlying assumption of cross-linking technologies is that RBPs require direct contact with mRNA binding partners to affect their stability, delivery or translation. However, mechanistically, this may not be correct, and the isolation of RNA immunoprecipitates by ESP does not require direct contact. For example, any given RBP may be required for the assembly or stability of ribonucleoprotein super-molecular complex granules with sequestered RNA species (49). Thus, although there is value in understanding the precise sites for RNA interaction (for example for FMRP (50)), this is not always necessary.
iv. *Parallel handling and rapid processing.* Prior characterization of the performance of the ESP devices has shown that specific but low-affinity reactions are preserved by rapid pull-through of magnetic beads through air or oil. This enhances the typical RIP-ChIP protocols by offering an expedited processing procedure, making the starting material less prone to degradation, and requiring less handling, which makes loss of key species during wash procedures less likely.

Note that the results of this assay are limited by the nature of the analysis of the complex components; here the results are restricted to probes that appeared on the micro-arrays, eliminating important regulatory non-mRNA species such as lncRNA, ncRNA or miRNAs. However, this technique is entirely compatible with other techniques such as RNA-Seq, or targeted arrays, which would reveal these species. It is also dependent upon the antibodies used for immunoprecipitation: for example, antibodies previously reported to identify CRD-BP/IGF2BP1 partners did not pass our validation criteria (7, 51). Furthermore, mass spectrometry is entirely compatible with ESP-RIP, allowing a full dissection of protein components of RBP particles.

### 4.3 Comparison of the results of ESP purification of CRD-BP binding RNAs with other studies

CRD-BP has been implicated in a variety of aspects of mRNA metabolism and expression, from the stabilization of mRNAs by blocking miRNA binding sites (52, 53), to the localization or translation of cognate proteins (54, 55). Functionally, this protein it is important to cell survival, cell migration and chemo-resistance (56–59), and this underlies the focus on determining its mechanism of action.

There are parallel data for several types of RIP analyses of CRD-BP in different cell types; the results of five studies, including this one, are shown in Table 1 (7, 36, 37, 60). Nielsen and colleagues exogenously expressed Flag-tagged CRD-BP to determine potential mRNA interactions in 293T cells; they found that CRD-BP associates with a considerable proportion of the total transcriptome (3%) in large 100-300 nm intracellular granules (36). The study started with 100× more cell lysate, and did not report total yields (Jonson et al., 2007). Overall, the results showed 352 specific mRNA species associated with CRD-BP, which is in the same order of magnitude as our study, where our study relied on the low endogenous expression in mouse mammary cells as the immunoprecipitation target. Furthermore, assay of mRNA binding partners identified by Jonson et al confirmed that these were also enriched in CRD-BP associated RNA populations from mammary epithelial cells in this study (Fig. 5C).

**Table 1.** Comparison of RNA associations for CRD-BP defined by different RIP techniques. The experimental conditions are listed that relate to approach, cell background and outcomes of each technique.

|  | **Method** | **Scale/**<br>**Cell type** | **CRD-BP**<br>**target** | **RNA cross-**<br>**link / RNA**<br>**labeling** | **Assay** | **Proportion of CRD-BP**<br>**retrieved for analysis;**<br>**number of RNA binding**<br>**partners** |
| --- | --- | --- | --- | --- | --- | --- |
| Jonson<br>et al 2007<br>(36) | RIP | 1.20 x 10<sup>8</sup><br>293T cells | Flag-tagged<br>Exogenous | None<br>None | Affymetrix<br>U133 Plus 2.0<br>array | ND<br>3% total mRNA in multi-<br>complex granules; 352<br>mRNAs<br>(>3x enriched, p<0.05) |
| Hafner et<br>al 2010<br>(5) | PAR-CLIP | > 10<sup>8</sup><br>293T cells | Flag/HA-tagged<br>Exogenous | UV Xlink<br>Labeling in<br>vivo with 4SU;<br>labeled in vitro<br>with <sup>32</sup>P | Solexa<br>sequencing of<br>cDNA library;<br>ID of<br>interacting<br>sequences | ND<br>56 mRNAs (+19 mRNAs<br>encoded by mitochondrial<br>genome) |
| Barnes et<br>al 2015<br>(60) | RIP | ~10<sup>6</sup><br>HeLa<br>cells | Flag/HA-tagged<br>mouse<br>sequence<br>Exogenous | None<br>None | Site directed<br>mutagenesis<br>of CRD-BP;<br>focus on three<br>binding<br>partners,<br>CD44, c-myc<br>and β-actin | ND<br>Not known |
| Conway<br>et al 2016<br>(7)<br>Lambert<br>et al 2014<br>(61) | eCLIP | 2 x 10<sup>7</sup><br>hESC<br>cells | Endogenous<br>Flag-tagged<br>Exogenous | UV Xlink<br>Labeling in<br>vivo with 4SU;<br>labeled in vitro<br>with <sup>32</sup>P | Comparison<br>of eCLIP and<br>RBNS hits | ND<br>Not known |
| This<br>study | RIP with<br>ESP | <10<sup>6</sup><br>293T or<br>mammary<br>epithelial<br>cells | Endogenous<br>Flag-tagged<br>Exogenous | None | Nimblegen<br>array | 50-90% IGF2BP1 protein<br>900 RNA species enriched<br>(p<0.05) |

This contrasts with the lack of consensus from the published CLIP studies of CRD-BP: thus, the data sets derived from PAR-CLIP / 293T cells (37) and eCLIP / human embryonic stem cells (7) showed little overlap between gene sets derived from either CLIP technology and those derived from RIP (Jonson et al or this study). Specifically, there were only 6 matching mRNAs for the large libraries identified as CRD-BP binding partners in 293T cells (36, 37), and 17 matches between human 293T cells and this study of mouse mammary epithelial cells (36). This is despite shared expression of many of the mRNA species identified as CRD-BP partners. CLIP cross-linking techniques include sophisticated statistical arguments to identify significant rates of association of specific RNA sequences, leading Hafner et al to identify 56 unique mRNAs from RIP isolates of 293T cells transfected with tagged CRD-BP; the catalog of mRNAs identified in RIP isolates of human embryonic stem cells were not specified (7). The latter study states there was no overlap between the eCLIP gene set and the genes identified by the RNA Bind-n-Seq technique described by Lambert et al (61), or between those gene lists and the mRNAs destabilized by knockdown of CRD-BP. The reasons for the lack of overlap between the parallel techniques used in these studies is not yet understood and could be important to resolve. Note that CRD-BP was originally isolated as the protein protecting c-Myc mRNA from degradation in K562 cells (16, 53, 62), but this association is not necessarily typical of other cell types, despite widespread expression of c-Myc.

We are intrigued by the enrichment of mRNAs for proteins destined to be imported into mitochondria (63) and for components of the glutathione metabolism pathway (27, 64). There is a well-established partnership between regulated RBP activity and mitochondrial mRNA translation and import (38). Mitochondrial function in turn is an important determinant of clonogenic survival and establishment, which are consistent markers of CRD-BP activity (65, 66). For example, the RBP CLUH (clustered mitochondria homologue), together with other RBPs, have been shown to regulate the expression of a mitochondrial protein network that become important under conditions of nutrient deprivation (67). Glutathione metabolism is key to the disposition of super-oxides, which are produced at high levels during chemotherapy (68). CRD-BP and its close paralogue, IGF2BP3, have beeen shown to enhance drug resistance for tumor cells (69, 70). However, we note that the specific gene ontology keywords differ for this study of mouse mammary epithelial cells and for the 293T cells described by Jonson et al (proteins involved in secretory pathway and endoplasmic reticulum-associated quality control, as well as ubiquitin-dependent metabolism), and the verification of the actual activity of CRD-BP awaits functional evaluation in cell types that show important CRD-BP dependent phenotypes.

## 5. Conclusions

We have demonstrated the application of a simple enhanced RNA immunoprecipitation procedure that enables parallel processing with highly efficient retrieval of specific RBP-associated mRNAs. This procedure can be applied to limiting amounts of experimental materials. This increases the currently available technologies that can be applied to this field of research and offers the opportunity to evaluate alterations of RBP-associated species under numerous experimental conditions, that is not feasible by other means. Our preliminary analysis of the RBP, CRD-BP/IMP1, in cultured mouse mammary epithelial cells reveals intriguing links to functional components of cancer cell metabolism.

## Acknowledgments

Our appreciation to Celia Bisbach and Josh Martin for their technical assistance, and to Gilson Inc (Madison WI) for their gift of the EXTRACTMAN device. SAF was supported by the Kuwait Foundation for the Advancement of Sciences under project 2013-6302-03. This project was also supported by a DOD Scholar Award W81XWH-06-1-0491 (CMA), NINDS R21NS095187 (AR), NCI R01CA186134 (DJB) and NIAMS-RO1AR063361 and NCI-RO1CA121851 (to VSS). This work was also supported in part by pilot funding from the NIH/NCI P30 CA014520--UW Comprehensive Cancer Center Support.

More information on the application of these SLIDE devices is presented in video form at https://www.gilson.com/extractman-starter-kit.html.

## Author contributions

CMA, SMB and SAF designed research; SAF performed research; AR, SMB and DJB contributed devices, analytic tools and bioinformatic expertise; AR, CMA, VSS and SAF analyzed data; CMA and SAF wrote the paper.

## Abbreviations

ESP™: exclusion-based sample preparation
IFAST™: Immiscible Filtration Assisted by Surface Tension
SLIDE™: Sliding Lid for Immobilized Droplet Extractions
RBP: RNA binding protein
RIP: RNA immunoprecipitation
RNP: ribonucleoprotein
PMPs: paramagnetic particles
CRD-BP: coding region determinant-binding protein
IMP1: insulin-like growth factor-2 mRNA binding protein 1
PAR-CLIP: photoactivatable-ribonucleoside-enhanced cross-linking and immunoprecipitation

